# Chitosan primes plant defence mechanisms against *Botrytis cinerea*, including expression of Avr9/Cf-9 rapidly-elicited genes

**DOI:** 10.1101/2020.04.01.019513

**Authors:** Daniel De Vega, Nicola Holden, Pete E Hedley, Jenny Morris, Estrella Luna, Adrian Newton

## Abstract

Current crop protection strategies against the fungal pathogen *Botrytis cinerea* rely on a combination of conventional fungicides and host genetic resistance. However, due to pathogen evolution and legislation in the use of fungicides, these strategies are not sufficient to protect plants against this pathogen. Defence elicitors can stimulate plant defence mechanisms through a phenomenon known as priming. Priming results in a faster and/or stronger expression of resistance upon pathogen recognition by the host. This work aims to study priming of a commercial formulation of the elicitor chitosan. Treatments with chitosan result in induced resistance in solanaceous and brassicaceous plants. In tomato plants, enhanced resistance has been linked with priming of callose deposition and accumulation of the plant hormone jasmonic acid (JA). Large-scale transcriptomic analysis revealed that chitosan primes gene expression at early time-points after infection. In addition, two novel tomato genes with a characteristic priming profile were identified, Avr9/Cf-9 rapidly-elicited protein 75 (*ACRE75)* and 180 *(ACRE180)*. Transient and stable overexpression of *ACRE75, ACRE180* and their *Nicotiana benthamiana* homologs, revealed that they are positive regulators of plant resistance against *B. cinerea*. This provides valuable information in the search for strategies to protect Solanaceae plants against *B. cinerea*.

## Introduction

Crop yield losses of 20-40% of total agriculture productivity can be attributed to pests and diseases (Oerke, 2006, Savary et al., 2012). Of these threats, the pathogen *Botrytis cinerea* causes annual losses of $10-$100 billion, as it reduces crop yield before harvest or leads to waste and spoilage post-harvest. It is the causative agent of grey mould disease in tomato and many other economically important crops, such as pepper, aubergine, grape, lettuce and raspberry. *B. cinerea* is a fungal generalist (broad-host range) and considered to be a model necrotrophic pathogen (Williamson et al., 2007). Effective control include the use of conventional crop protectants (e.g. fungicides) and resistant varieties as well as sanitation and environmental control. However, rapid pathogen evolution can result in the loss of efficacy of resistance sources and fungicides (Pappas, 1997, Williamson et al., 2007). In addition, the use of pesticides is strictly limited by European regulations due to human health and environment risk and hazard assessment changes. New alternative strategies are therefore needed. Exploiting the plant’s defence system to provide protection against these threats has emerged as a potential strategy against pathogen infection and disease (Luna, 2016).

Plant endogenous defences is activated by elicitor molecules resulting in induced resistance (IR) (Mauch-Mani et al., 2017), since they are able to mimic pathogen-inducible defence mechanisms (Aranega-Bou et al., 2014). Induced resistance works via two different mechanisms: direct activation of systemic plant defences after signal recognition and; priming, a mechanism that initiates a wide reprogramming of plant processes, considered to be an adaptive component of induced resistance (Mauch-Mani et al., 2017). Priming has been demonstrated to be the most cost-effective mechanism of induced resistance in terms of plant development as there is no direct relocation of plant resources from growth to defence until it is necessary (van Hulten et al., 2006). Studies have already shown that low elicitor doses can enhance resistance to pests without interfering with crop production (Redman et al., 2001). Elicitor-induced priming has been demonstrated to last from a few days (Conrath et al., 2006) to weeks (Worrall et al., 2012) after treatment and even through subsequent generations (Ramírez-Carrasco et al., 2017, Slaughter et al., 2012).

Priming can have multiple effects on plant defences, which vary depending on the type of plant-pathogen interaction. Defence priming enables the plant to fine-tune immunity responses through enhancement of the initial defences. These is achieved through different mechanisms that act at specific defence layers (Mauch-Mani et al., 2017). For instance, cell-wall fortification and effective production of reactive oxygen species (ROS) has been used as a marker for the expression of priming responses. Hexanoic acid (Hx) primes cell-wall defences through callose deposition and redox processes in tomato cultivars against *B*.*cinerea* (Aranega-Bou et al., 2014). In *Arabidopsis thaliana*, BABA and benzothiadiazole (BTH)-induced priming is also based on an increase in callose deposition (Kohler et al., 2002, Ton et al., 2005). Priming also results in transcriptomic changes. Gene expression analysis of *A. thaliana* after BABA treatment was used to identify a transient accumulation of SA-dependent transcripts, including that of *NPR1*, which provides resistance against *Pseudomonas syringae* (Zimmerli et al., 2000). Changes in metabolite accumulation have been shown to mark priming of defence also. For instance, defence hormone profiling has shown that accumulation of JA and JA-derivatives mediates priming of mycorrhizal fungi (Pozo et al., 2015). Moreover, untargeted metabolomic analysis have identified different compounds, including kaempferol (Król et al., 2015), quercetin, and indole 3 carboxylic acid (I3CA) (Gamir et al., 2014), that drive priming responses.

Several elicitors have been described to induce resistance mechanisms in tomato against *B. cinerea*. For instance, BABA has been demonstrated to provide long-lasting induced resistance against *B. cinerea* in leaves (Luna et al., 2016) and in fruit (Wilkinson et al., 2018). In addition, the plant defence hormone JA has also been linked to short-term and long-term induced resistance in tomato against *B. cinerea* (Luna et al., 2016, Worrall et al., 2012). To date, however, few studies have investigated elicitor-induced priming in tomato against *B. cinerea*. One of them showed that Hx-induced priming is based on callose deposition, the expression of tomato antimicrobial genes (e.g. protease inhibitor and endochitinase genes), and the fine-tuning of redox processes (Aranega-Bou et al., 2014, Finiti et al., 2014). Therefore, evidence is building in tomato, that induced resistance against *B. cinerea* can be based on priming also.

In this study we investigated whether the chitin de-acetylated derivative, chitosan, triggers priming of defence in tomato against *B. cinerea*. Chitosan as a plant protection product is considered ‘generally recognised as safe’ (Raafat and Sahl, 2009) that is effective in protecting strawberry, tomato and grape against *B. cinerea* (Muñoz and Moret, 2010, Romanazzi et al., 2013). Different studies have shown that its effect on crop protection results from induction of defence mechanisms (Sathiyabama et al., 2014) and direct antimicrobial activity (Goy et al., 2009). However, treatments with chitosan require infiltration into the leaves to trigger a robust effect (Scalschi et al., 2015) making it an unsuitable method of application in large-scale experiments or studies that take into consideration first barrier defence strategies. Here, we have addressed whether treatment with a water-soluble formulation of chitosan results in induced resistance phenotypes and in priming of cell wall defence and defence hormone accumulation. In addition, whole-scale transcriptome analysis was performed to identify candidate genes that are driving expression of priming. Our findings, together with the outlined characteristics of chitosan, make this substance a suitable candidate for extensive application as a component of Integrated Pests (and disease) Management (IPM) for the protection of crops against fungal pathogens.

## Material and Methods

### Plant material and growth conditions

Tomato cv. Money-maker seeds were used in the described experiments. Unless otherwise specified, seeds were placed into propagator trays containing Bulrush peat (Bulrush pesticide-free black peat, low nutrient and low fertilizer mix) and a top layer of vermiculite and left at 20 °C until germination. Germinated seeds were transplanted to individual pots containing Bulrush soil (pesticide-free compost mix and nutrient and fertilizer rich) in a growth cabinet for 16h - 8h / day-night and 23°C / 20°C cycle C at ∼150 μE m^−2^ s^−1^ at ∼ 60% relative humidity (RH). *Nicotiana benthamiana* seeds were cultivated in a similar manner specified for tomato for 16h - 8h/ day-night cycle; 26°C / 22°C at ∼150 μE m^−2^ s^−1^ at ∼ 60% relative humidity (RH). Aubergine (*Solanum melongena*) cv. Black Beauty seeds were placed into propagators containing Bulrush peat and a layer of vermiculite on the top and incubated at 20°C for 1-2 weeks until germination. Seedlings were then transplanted to individual pots containing Bulrush soil and grown and cultivated as for tomato. *Arabidopsis thaliana* (hereafter referred to as Arabidopsis) Columbia-0 (Col-0) and transgenic lines were grown in a soil mixture of 2/3 Levington M3 soil and 1/3 sand for 8h - 16h/day - night and 21°C / 18°C cycle at ∼150 μE m^−2^ s^−1^ at ∼ 60% RH. Ten-day-old plants were transplanted to individual pots.

### Chemical treatment

All experiments were performed using a commercial, water-soluble chitosan formulation, known as ChitoPlant (ChiPro GmbH, Bremen, Germany) (Romanazzi et al., 2013, Younes et al., 2014). ChitoPlant, referred to as chitosan latterly, was freshly prepared in water to the specific concentrations (please see figure legends for details). Treatments were performed by foliar spraying of chitosan solution (with 0.01% Tween20) directly onto leaves.

### *Botrytis cinerea* cultivation, infection and scoring

*B. cinerea* R16 (Faretra and Pollastro, 1991) was used in all experiments and was kindly provided by Dr Mike Roberts (Lancaster University). Cultivation of the fungus and infection of tomato-based experiments were performed as described (Luna et al., 2016). For *N. benthamiana*, 2-3 detached leaves were inoculated with 6µl inoculum solution containing 2 × 10^4^ spores/ml of *B. cinerea*. Infected leaves were kept at 100% RH by sealing the trays and placed in the dark before disease assessment. Arabidopsis infections were performed as previously described (La Camera et al., 2011) with a few modifications. Leaves were inoculated with 5µl inoculum solution containing ½ strength of potato dextrose broth (PDB – Difco at 12 g/l) and 5 × 10^5^ spores/ml. Infected Arabidopsis plants were put in a sealed tray at 100% RH and moved back to the growth cabinet. Infections of *S. melongena* plants were performed by drop inoculating detached leaves with a spore solution of *B. cinerea* containing 2 × 10^4^ spores/ ml. For all pathosystems, disease was scored by measuring lesion diameters with an electronic calliper (0.1 mm resolution) on different days post infection.

### Plant growth analysis

Relative growth rate (RGR) was used to analyse tomato growth after chitosan treatment as described (Luna et al., 2016). Growth analysis of Arabidopsis plants was performed by measuring rosette perimeter using Photoshop CS5 (Vasseur et al., 2018).

### Callose deposition assays

For analysis of callose deposition after chitosan treatment, material from tomato and Arabidopsis plants with different concentrations of chitosan were collected 1 day after treatment (dat) and placed in 96% (v/v) ethanol in order to destain leaves. Aniline blue was used to stain callose deposits as described previously (Luna et al., 2011). Analysis of callose associated with the infection by *B. cinerea* in tomato leaves was performed as described (Rejeb et al., 2018) with some modifications. Briefly, infected tomato leaf samples were collected and placed in 96% (v/v) ethanol 1 day after infection with *B. cinerea* and allowed to destain. Destained material was hydrated with 0.07 M phosphate buffer (pH 9.0) for 30 min and then incubated for 15 min in 0.1% (w/v) aniline blue (Sigma-Aldrich) and 0.005% (w/v) fluorescent brightener (Sigma-Aldrich). Solutions were then replaced with 0.1% (w/v) aniline blue and incubated for 24h in the dark prior to microscopic analysis. All observations were performed using an UV-epifluorescence microscope (GXM-l2800 with GXCAM HiChrome-MET camera). Callose was quantified from digital photographs by the number of yellow pixels (callose intensity). Infection-associated callose was scored and analysed in a similar way but callose intensity was expressed relative to fungal lesion diameters. Image analyses were performed with Photoshop CS5 and ImageJ.

### Chitosan antifungal activity *in vitro* assay

*B. cinerea* mycelial growth assessment was performed using Potato Dextrose Agar (PDA) as culture media with different concentrations of chitosan (1%, 0.1%, 0.01% w/v). PDA was autoclaved and then chitosan and the fungicide Switch (as positive fungicide control) (1%, 0.1%, 0.01% w/v) were added directly to PDA as it cooled. One 5 mm diameter agar plugs of actively growing *B. cinerea* mycelium was added per plate. Five plates per treatment were sealed with parafilm and then incubated under controlled conditions (darkness and 24°C). After 4 days, the mean growth of the fungus was determined by measuring two perpendicular diameters and calculating the mean diameter.

### High-Pressure Liquid Chromatography (HPLC) - Mass Spectrometry (MS)

Healthy and infected tomato leaf tissues were harvested in liquid nitrogen and subsequently freeze-dried for 3 days. Freeze-dried samples were ground in 15 mL Falcon tubes containing a tungsten ball in a bead beater. Ten mg of each sample was used for hormone extraction. Sample extraction, HPLC-MS quantitative analysis of plant hormones and data analysis were performed as described (Forcat et al., 2008). Accurate quantification of ABA, SA and JA used the deuterated internal standards added during sample extraction (Forcat et al., 2008) and concentrations were calculated using standard concentration curves. Relative accumulation of jasmonic acid-isoleucine (JA-Ile) were obtained by calculations of % peak areas among samples.

### Transcriptome analysis

Four conditions were analysed using microarrays: **(i)** ddH_2_O-treated and non-infected plants (Water + Mock); **(ii)** Chitosan-treated and non-infected plants (Chitosan + Mock); **(iii)** ddH_2_O-treated and *B. cinerea*-infected plants (Water + *B. cinerea*); **(iv)** Chitosan-treated and *B. cinerea*-infected plants (Chitosan + *B. cinerea*). Inoculations were performed four days after treatment (dat) with chitosan, and leaf discs from four independent plants (biological replicates) per treatment were sampled at 6, 9 and 12 h post-inoculation (hpi) with mock or *B. cinerea* spores. Total RNA was extracted with an RNeasy Plant Mini Kit (Qiagen) as recommended. A custom 60-mer oligonucleotide microarray was designed using eArray (https://earray.chem.agilent.com/earray/; A-MTAB-667 and E-MTAB-8868; www.ebi.ac.uk/arrayexpress/) from predicted transcripts (34,616 in total) of the *S. lycopersicum* (ITAG 2.3) genome. Experimental design is detailed at E-MTAB-8868; www.ebi.ac.uk/arrayexpress/. Two-channel microarray processing was utilised, according to the Low Input Quick Amp Labelling Protocol v. 6.5 (Agilent). Microarray images were imported into Feature Extraction software (v. 10.7.3.1; Agilent) and data extracted using default parameters. Data were subsequently imported into Genespring software (v. 7.3; Agilent) for subsequent pre-processing and statistical analysis. Following Lowess normalisation, data were re-imported as single-colour data. Data were filtered to remove probes that did not have detectable signal in at least 3 replicates, leaving 22,381 probes for statistical analysis.

Analysis of Variance (2-way ANOVA; p-value ≤0.01, Benjamini-Hochberg false discovery rate correction) was used to identify differentially expressed genes (DEGs) for the factors ‘Treatment’ (3,713 DEGs), ‘Time’ (6,920), and ‘Treatment-Time interaction’ (186). Subsequently, pairwise Student’s T-tests were performed (Volcano plots: P-value ≤0.05, 2-fold cut-off) on the global set of 8,471 DEGs for each of the three test treatments (Chitosan + Mock, Water + Mock and Chitosan + *B. cinerea*) compared to control (Water + Mock) at each time point. Venn diagrams were used at each time point to identify common and specific DEGs.

### Panther gene ontology (GO) term enrichment analysis

Panther software (Thomas et al., 2003) was used to visualise DEG products in the context of biological pathways and/or molecular functions, using default settings. Functional enrichment analysis was performed using DEG lists for Chitosan + *B. cinerea* and Water + *B. cinerea* treatments at 6 hpi. ‘Biological processes’ and ‘molecular functions’ were selected using PANTHER Overrepresentation Test (release 20170413) against *S. lycopersicum* (all genes in database) and Bonferroni correction for multiple testing.

### DEG transcript co-expression analysis

Two-way ANOVA was performed on the filtered microarray dataset at increased stringency (p-value ≤0.01, Bonferroni false discovery rate correction) to identify 1,722 highly-significant DEGs. Pearson’s correlation was used with default settings in Genespring (v 7.3) to generate a heatmap to help identify co-expressed transcripts (Figure 3b).

**Figure 1.**
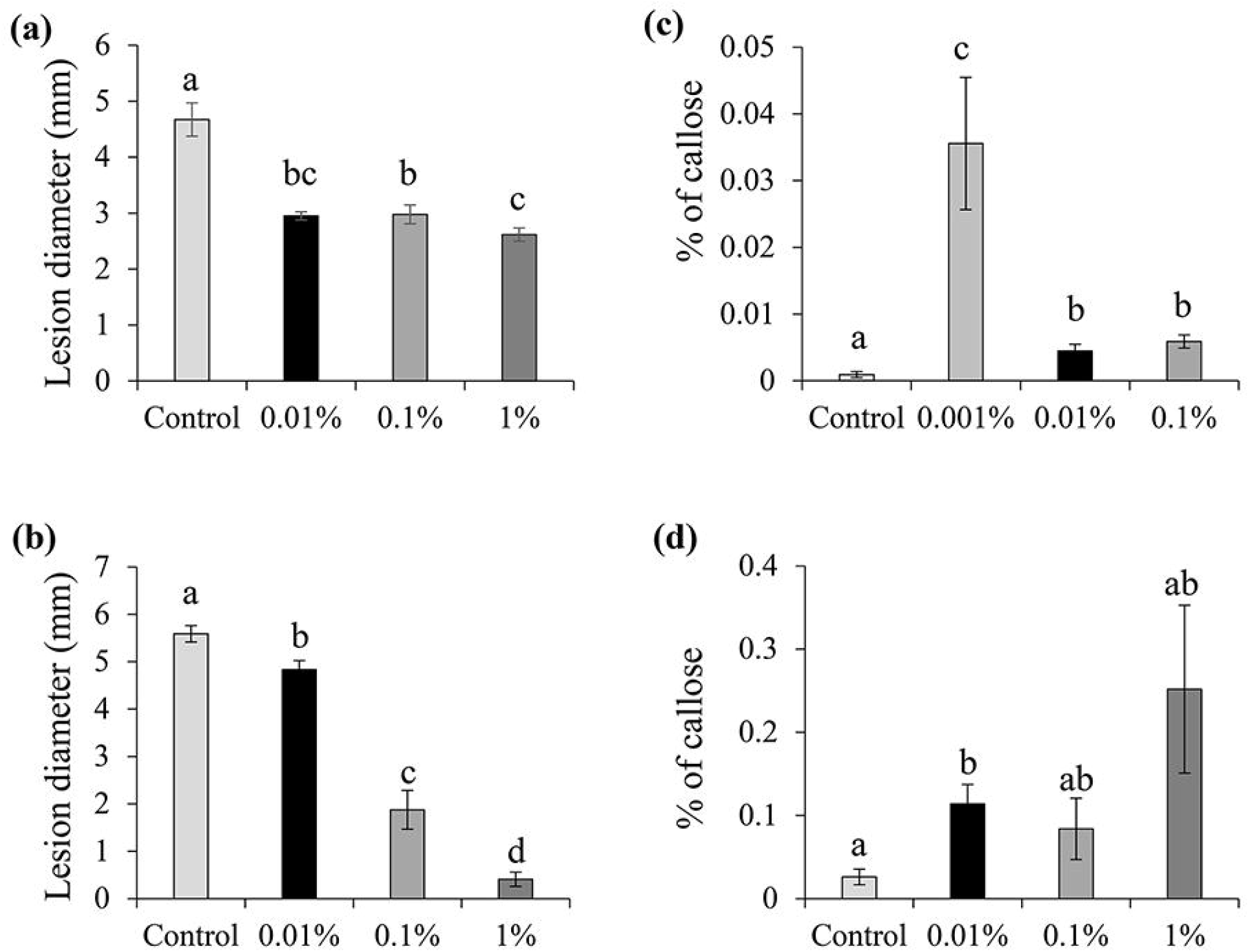
Characterisation of chitosan-induced resistance in tomato and Arabidopsis. (a) Disease lesions in tomato and (b) in Arabidopsis at 3 days post inoculation. Values represent means ± SEM (n=4-10). (c) Callose deposition triggered by chitosan treatment in tomato and (d) in Arabidopsis 1 day post treatment. Values represent means ± SEM (n=8-10) of % of callose per leaf area. Different letters indicate statistically significant differences among treatments (Least Significant Differences for graph a and Dunnett T3 Post-Hoc test for graphs b, c and d, α=0.05).

**Figure 2.**
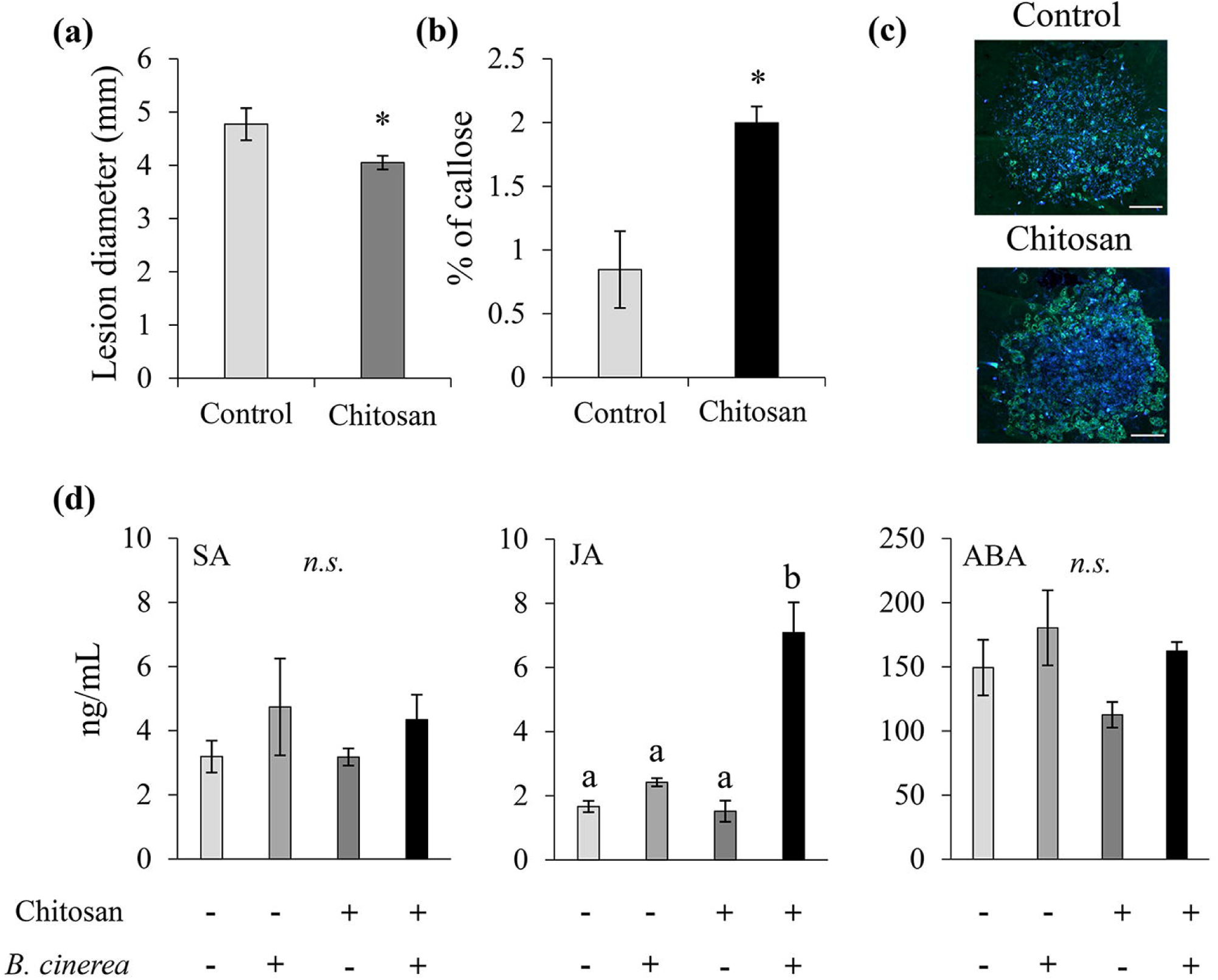
Chitosan-induced resistance is based on priming. **(a)** Disease lesions in tomato at 3 days post inoculation (dpi) 2 weeks after treatment with water (Control) or 1% chitosan. **(b)** Values represent means ± SEM (n=8). Asterisk indicates statistically significant differences among treatments (Student’s T. test, α=0.05). **(b)** Percentage of callose deposited at the infection site in water (Control) and chitosan (0.01%)-treated plants compared to the fungal lesion diameter at 1 day after infection with *B. cinerea*. Values represent means ± SEM (n=4). Asterisk indicates statistically significant differences among treatments (Student’s T. test, α=0.05). **(c)** Representative pictures of chitosan-induced priming of callose at the infection site. Blue colours correspond to fungal growth whereas yellow colours correspond with the callose deposition at the infection site. Scale bars= 0.5 mm. **(d)** Mass-spectrometry quantification (ng/mL) of Salicylic acid (SA), Jasmonic acid (JA) and Abscisic acid (ABA) at 24h post infection. Values represent means ± SEM (n=4). letters indicates statistically significant differences among treatments (Least Significant Differences, α=0.05, n.s. = not significant).

**Figure 3.**
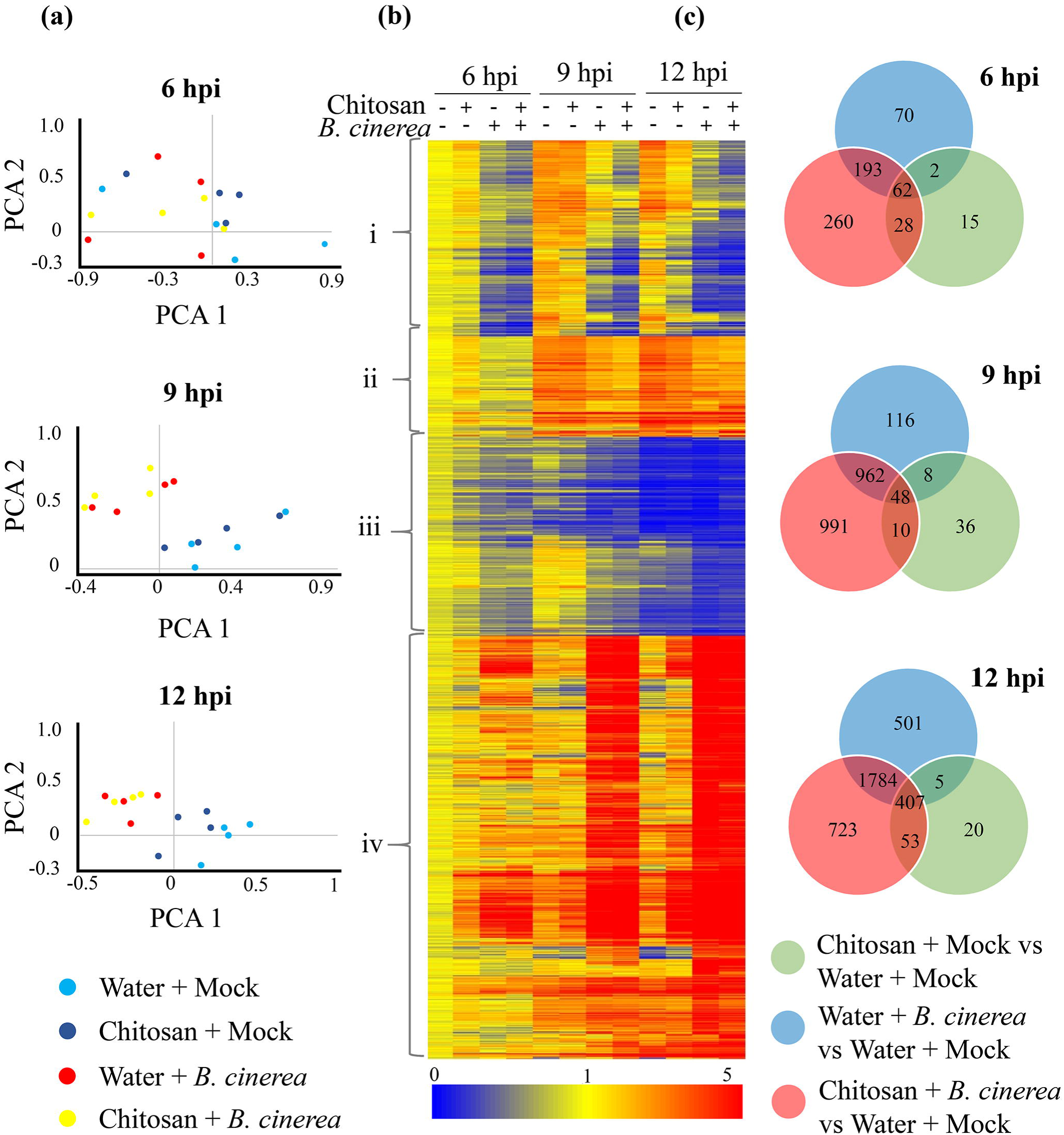
Transcriptome analysis of chitosan-induced resistance against *Botrytis cinerea*. **(a)** Principal component analysis (PCA) of whole transcriptional microarray at 6, 9 and 12 hour post infection (hpi). **(b)** Heatmap of differentially expressed genes (2-way ANOVA *p* < 0.01, Bonferoni), clustered by expression. Profiles are shown +/- treatment with chitosan and infection with *B. cinerea* at 6, 9, 12 hpi logarithmic scale fold induced (red) or repressed (blue) compared to Water + Mock. Hierarchical clusters on expression profile are broadly classed as i, ii, iii or iv. **(c)** Venn diagram of statistically significant data set (2-way ANOVA *p* < 0.01, Benjamin–Hochberg) of differentially expressed genes. Pairwise Student’s T-test comparisons were performed (Volcano plots: *p* < 0.05, 2-fold cut-off) for the three test treatments (Chitosan + Mock, Water + *B. cinerea* and Chitosan + *B. cinerea*) compared to control treatment (Water + Mock) at 6, 9 and 12 hpi.

### Gene expression analysis

Validation of *S. lycopersicum* transcriptomic analysis was performed by qRT-PCR of nine candidate differentially expressed genes (DEGs), comparing gene expression values with microarray. RNA samples were DNAse-treated with TurboDnase (ThermoFisher) and complementary DNA (cDNA) was synthesized from 2.5 µg total RNA using Superscript III reverse transcriptase (Invitrogen) as recommended with random hexamer/oligo dT primers. RT-qPCR reactions were performed with specific *S. lycopersicum* oligonucleotide primers (Table S4) purchased from Sigma-Aldrich. Gene primers and probes were designed using Universal Probe Library (UPL) assay design centre (Roche Diagnostics Ltd.). RT-qPCR was performed using FastStart Universal Probe Master Mix (Roche) and expression was calculated against two reference genes (*SlActin-like* and *SlUbiquitin*) using the Pfaffl method (Pfaffl, 2001).

### Gene cloning

Orthologues of *SlACRE75* and *SlACRE180* were obtained from CDS and protein sequences BLAST analysis against Arabidopsis genome (TAIR10) for Arabidopsis sequences, or a reciprocal best BLAST hits (RBH) (Ward and Moreno-Hagelsieb, 2014) test was performed (Sol Genomics Network) for *N. benthamiana*, termed *NbACRE75* and *NbACRE180*, respectively. Best CDS and protein hits were identified, being Niben101Scf03108g12002.1 and Niben101Scf12017g01005.1 for SlACRE75 and SlACRE180 respectively; termed NbACRE75 and NbACRE180 onwards. Flanking the (i) *SlACRE75*, (ii) *SlACRE180*, (iii) *NbACRE180* and (iv) *NbACRE75* coding sequences (CDS), Gateway® cloning was used to design and produce overexpression constructs for the gene candidates for a N-terminal GFP:ACRE fusion protein per insert (Reece-Hoyes and Walhout, 2018). Briefly, pUC57 plasmids containing *SlACRE75, SlACRE180, NbACRE75* and *NbACRE180* coding sequences were chemically synthesized by GenScript. For *SlACRE75, SlACRE180, NbACRE75* and *NbACRE180*, cDNAs from pUC57 entry vector were transformed by electroporation into *Escherichia coli* strain DH10B and transferred by a recombinant LR reaction of Gateway cloning (Clonase II enzyme mix Kit, Thermo Fisher) into pB7WGF2 (Karimi et al., 2002).

### Transient expression in *Nicotiana benthamiana*

*Agrobacterium tumefaciens*, strain GV3103, carrying plasmids with expression constructs (i) pB7WGF2:35S:GFP:SlACRE75; (ii) pB7WGF2:35S:GFP:SlACRE180; (iii) pB7WGF2:35S:GFP:NbACRE180; (iv) pB7WGF2:35S:GFP:NbACRE75 and; (v) pB7WGF2:35S:GFP (empty vector), were grown in YEP medium (containing 50 µg/ ml rifampicin, 100 µg/ ml spectinomycin, and 25 µg/ ml gentamicin) for 24 h with continuous shaking at 28°C. Overnight cultures were collected by centrifugation, resuspended in Agromix/infiltration buffer (10 mM MgCl_2_ : 10 mM MES) and 200 µM acetosyringone (pH 5.7) and diluted to a final volume of 20 ml at OD_600_ of 0.1. Cultures were infiltrated into leaves of 4-week-old *N. benthamiana* plants using 1 ml needleless syringes. One day after agroinfiltration, 1-2 leaves per plant were excised for *B. cinerea* infection assays (as described above). These experiments were repeated once.

### Confocal microscopy analysis

For the analysis of the subcellular localization, *A. tumefaciens* GV3101 carrying plasmids with expression constructs were co-infiltrated with pFlub vector (RFP-peroxisome tagged marker) into leaves of 4-week-old *N. benthamiana* CB157 (nucleus mRFP marker) and CB172 (ER mRFP marker) reporter lines using 1 ml needleless syringes. Two days after infiltration, leaves were excised and prepared for confocal microscopy. GFP and mRFP fluorescence was examined under Nikon A1R confocal microscope with a water-dipping objective, Nikon X 40/ 1.0W. GFP was excited at 488 nm from an argon laser and its emissions were detected between 500 and 530 nm. mRFP was excited at 561 nm from a diode laser, and its emissions were collected between 600 and 630 nm.

### Western blot analysis

Leaves from *N. benthamiana* leaves infiltrated with *A. tumefaciens* GV3101 carrying plasmids with expression constructs were excised, ground and proteins extracted as previously described (Gilroy et al., 2011, Yang et al., 2016). Western blotting was performed as previously described (Qin et al., 2018). Detection of GFP was performed using a polyclonal rabbit anti-GFP antibody (1:4,000 dilution) and secondary anti-mouse antibody (IG HRP 1:10,000) according to the manufacturer’s instructions. ECL development kit (Amersham) detection was used according to the manufacturer’s instructions.

### Transformation of *Arabidopsis thaliana* stable overexpression transgenic lines

Arabidopsis overexpression plants were transformed using *A. tumefaciens* GV3101 carrying plasmids with expression constructs using the flower dipping method (Clough and Bent, 1998). Selection of Arabidopsis transformants and homozygous lines selection were performed as described (Luna et al., 2014). Resistance was tested against *B. cinerea* as described before. Two independent homozygous overexpression lines were obtained per construct.

### Pathosystem statistics

Statistical analysis of induced resistance and growth phenotypes were performed as described (Luna et al., 2016). Data analysis was performed using SPSS Statistics 23 and GenStat® 18th Edition (VSN International, Hemel Hempstead, UK). Statistical analysis of resistance phenotypes in Arabidopsis overexpression lines was done by ANOVA with ‘construct’ as a single treatment factor at 10 levels: Col-0 (wild-type treatment); two empty vector lines ‘EV 3.1’ and ‘EV 4.1’; ‘SlACRE75 1.1’ and ‘SlACRE75 2.1’; ‘SlACRE180 1.2’ and ‘SlACRE180 3.1; ‘NbACRE180 1.1’ and ‘NbACRE180 2.1’; and ‘NbACRE75 1.1’. The replicate units were individual plants of which there were 8-16 for each construct. Measurements of four lesions were recorded for each plant. Random effects were modelled as plant + plant × lesion to capture the plant-to-plant and within-plant variation. As part of the ANOVA, specific planned (non-orthogonal) contrasts were included to test for significant differences between the mean for each construct line compared to Col-0.

## Results

### Identification and characterisation of a novel chitosan formulation in its ability to induce resistance against *Botrytis cinerea*

We tested the water-soluble chitosan-based commercial formulation ChitoPlant, from hereafter termed chitosan, in its capacity to induce resistance against the fungal pathogen *B. cinerea*. Treatments of chitosan demonstrated that this elicitor successfully triggers resistance in tomato (Figure 1a), Arabidopsis (Figure 1b) and aubergine (Fig S1) against *B. cinerea*. In tomato, chitosan significantly decreased necrotic lesion size in all concentrations compared with control plants (Fig 1a). The resistance phenotype induced by chitosan had a dose-dependent effect at the two high concentrations (1% and 0.1%), however, the lowest concentration (0.01%) induced a level of resistance in between 0.1% and 1% treatments. In Arabidopsis, chitosan treatment resulted in induced resistance in a concentration-dependent manner, with 1% having the strongest effect (Fig 1b). In aubergine, chitosan treatment resulted in differences in lesion diameter in all concentrations compared to water-treated control plants (Figure S1), however, post-hoc analysis demonstrated that 0.1% was the most effective concentration.

We then tested whether chitosan induces callose deposition in a similar manner to other chitosan formulations (Luna et al., 2011). Plants were treated with increasing concentrations of chitosan one day before aniline blue staining. In both plant species, treatments with chitosan resulted in a direct induction of callose. The lowest concentrations of 0.001% and 0.01% in tomato and Arabidopsis, respectively, triggered the strongest effect (Fig 1 c and d).

To determine any antifungal effect of chitosan, different concentrations were tested on *B. cinerea* hyphal growth *in vitro* and compared to different concentrations of the fungicide Switch (Syngenta). Whereas all concentrations of Switch arrested pathogen growth, only 0.1% concentration of chitosan or higher had an antifungal effect (Fig S2). However, the lowest concentration of chitosan tested (0.01 %) had no antifungal effect compared to the control. This shows a concentration threshold for chitosan-direct antifungal activity against *B. cinerea*. Since 0.01% chitosan had no antifungal effect, but reduced *B. cinerea* lesions and induced callose formation, this concentration was selected for more in-depth analysis.

### Analysis of priming mechanisms marking chitosan-induced resistance

We tested whether induced resistance triggered by chitosan is mediated by priming mechanisms through the assessment of its capacity to induced long-lasting resistance in distal parts of the plants. Treatments with 1% chitosan induced long-lasting resistance against *B. cinerea* of up to 2 weeks after initial treatment of tomato plants (Fig 2a).

In order to assess whether treatments with chitosan directly affects plant development, we tested plant growth one week after treatment with 1% chitosan. These experiments revealed that chitosan treatment triggers a statistically significant growth promotion, therefore indicating that induced resistance by chitosan does not negatively impact plant development (Fig S3a).

To study whether chitosan induced resistance (IR) was based on known mechanisms of priming, callose and hormone profiling analysis were performed after subsequent infection. Treatment with chitosan resulted in the accumulation of approximately twice the callose deposited at the site of attack compared to plants treated with water (Fig 2b,c). In addition, mass spectrometry profiling of defence-dependent hormones demonstrated that chitosan-induced resistance is mediated specifically by accumulation of jasmonic acid (JA) (Fig 2c) and its amino acid conjugate JA-isoleucine (JA-Ile, Fig S3b). In contrast, no other impacts were found in the concentration of other defence hormones such as salicylic acid (SA) and abscisic acid (ABA) (Fig 2c). Thus, chitosan-IR is based on priming of callose at the infection site and accumulation of JA and its conjugate JA-Ile.

### Transcriptional analysis of chitosan-induced resistance

Priming of gene expression normally follows a characteristic pattern: differential expression is low, transient or often non-detectable after treatment with the elicitor only (i.e. Chitosan + Mock) and enhanced differential expression occurs upon subsequent infection (i.e. Chitosan + *B. cinerea*) compared to infected plants that were not pretreated with the chemical (i.e. Water + *B. cinerea*) (Conrath et al., 2006, Martinez-Medina et al., 2016). Importantly, the expression kinetics are also key points for the establishment of priming. To further determine the priming basis of chitosan-induced resistance, we performed whole transcriptomic analysis at 6, 9 and 12 hours post infection (hpi) with *B. cinerea*. These time points were selected as they cover the early, non-symptomatic start of the *B. cinerea* infection process. Unsupervised data analysis was first performed to observe global changes in the experiment. For this, we did a 2D principal component analysis (PCA) at different hours post infection. This analysis shows that chitosan treatments did not trigger major changes in transcription, however, it was the infection with *B. cinerea* which greatly impacts the experiment (Fig 3a). Moreover, whereas separation can be observed between Mock- and *B. cinerea*-infected replicates at 9 and 12 hpi, no obvious differences could be seen at the early time point of 6 hpi.

Genes with similar expression profiles were grouped, resulting in the identification of 1,722 differentially-expressed genes (DEGs) across all three treatments and time points. Hierarchical clustering separated the genes into four crude groups when compared to the Water + Mock treatment at the first time point (6 hpi, Fig 3b): Cluster *i* consists of genes that were repressed by *B. cinerea* infection; cluster *ii* represents genes induced by chitosan treatment only; cluster *iii* includes genes repressed by *B. cinerea* infection and by treatment of chitosan at the later time points; cluster *iv* consists of genes induced by *B. cinerea* infection and by treatment of chitosan only (Fig 3b). Overall patterns aligned with the previous finding that infection with *B. cinerea* had a large-scale, more extensive and differential response on tomato transcription compared to treatment with chitosan (Fig 3a). Moreover, the analysis demonstrates that application of chitosan results in a higher number of genes repressed than induced, with the exception of some highly induced genes in cluster *iv*. Distinct differences were evident between treatment with chitosan compared to infection with *B. cinerea*, e.g. a large group of genes in cluster *iv* differentially induced by *B. cinerea* at 9 and 12 h, as well as a large group of genes repressed by the pathogen in cluster *i*. This indicates that chitosan works as a priming agent that does not directly trigger major effects in gene transcription.

To study the different signalling pathways and specific genes responsible for priming of chitosan against *B. cinerea*, a two-way ANOVA identified 8,471 differentially expressed genes (DEGs) among all three treatments and time points. This global list of DEGs was subsequently used for focussed pairwise analysis to identify transcripts changing between treatments at each time-point. Venn diagrams demonstrates that the effect of chitosan on its own did not trigger major changes in gene transcription: only 15, 36 and 20 genes were differentially expressed in Chitosan + Mock vs Water + Mock treatments at 6, 9 and 12 hpi, respectively (Fig 3c). However, the effect of chitosan was much more pronounced after plants had been infected with *B. cinerea*. This combination resulted in the differential expression of 543, 2,011 and 2,967 genes at 6, 9 and 12 hpi, respectively, of which 260, 991 and 723 DEGs were induced only in the chitosan *B. cinerea* treatment (Fig 3c). In comparison, Water + *B. cinerea* treatments displayed differential expression of 327, 1,134, and 2,697 genes at 6, 9 and 12 hpi, respectively, of which 70, 116 and 501 DEGs were specific to the Water + *B. cinerea* treatment (Fig 3c). These results demonstrate that there is a subset of genes potentially responsible for chitosan-induced priming for a faster and more robust response against *B. cinerea*.

To further identify early-acting signalling pathways and genes involved in chitosan-induced priming, further analyses were performed on genes corresponding to the 260 probes differentially expressed only in the Chitosan + *B. cinerea* treatment at 6 hpi. Gene overrepresentation analysis was performed to identify biological processes and molecular functions of enriched genes. For biological processes, pathways such as response to stimulus, chemical and auxins were overrepresented (Table 1). Moreover, for molecular function, cysteine-type peptidase activity, transcription factor activity, sequence-specific DNA binding and nucleic acid binding transcription factor activity were enriched (Table 1).

**Table 1:** Biological processes and molecular functions of enriched genes.

| GO biological processes for Chitosan + B.cinerea at 6 hpi |  |  |  |  |  |  |
| --- | --- | --- | --- | --- | --- | --- |
|  | S. lycopersicum ref. # | Upload # | Expected | Fold Enrichment | +/- | P value <0.05 |
| response to auxin | 209 | 10 | 1.58 | 6.35 | + | 9.47E-03 |
| response to chemical | 916 | 20 | 6.91 | 2.9 | + | 4.55E-02 |
| response to stimulus | 2657 | 44 | 20.03 | 2.2 | + | 1.25E-03 |
| Unclassified | 17617 | 117 | 132.83 | 0.88 | - | 0.00E+00 |
| GO molecular functions for Chitosan + B.cinerea at 6 hpi |  |  |  |  |  |  |
|  | S. lycopersicum ref. # | Upload # | Expected | Fold Enrichment | +/- | P value <0.05 |
| cysteine-type peptidase activity | 271 | 11 | 2.04 | 5.38 | + | 1.23E-02 |
| transcription factor activity, sequence-specific DNA binding | 856 | 19 | 6.45 | 2.94 | + | 4.78E-02 |
| nucleic acid binding transcription factor activity | 856 | 19 | 6.45 | 2.94 | + | 4.78E-02 |
| Unclassified | 16331 | 109 | 123.14 | 0.89 | - | 0.00E+00 |

### Identification of genes primed by chitosan

To identify genes that could be involved in chitosan-induced resistance, gene expression profiles were scrutinized. First, qRT-PCR analysis of a subset of 9 genes was done to successfully validate the expression data of the microarray (Fig S4). Similar expression profiles were observed in the microarray and the qRT-PCR data, validating the data set. Priming profiles, i.e. subtle or non-detectable differential expression after chitosan treatment (i.e. Chitosan + Mock) and an increased differential expression after infection (i.e. Chitosan + *B. cinerea*) were identified. Transcripts of the earliest time point during the infection (6 hpi) were chosen to identify primed genes involved in early immune responses. Expression of the subset of 260 DEGs unique for Chitosan + *B. cinerea* treatment at 6 hpi (Fig 3c) were analysed over Water + Mock, Chitosan + Mock and water + *B. cinerea*. From the subset, 203 down-regulated (Table S1) and 57 genes were found to be up-regulated (Table S2). An over-representation test was performed to investigate gene ontology categories of the primed genes (Panther 14.0).

Among the 203 genes that were repressed during infection (Table S1), eleven transcripts were associated with cysteine-type peptidase activity. Other transcripts were grouped with photosynthesis, light harvesting in photosystem I activity. Moreover, several had a response to hormone activity; nine ethylene-responsive transcription factor and receptor genes were significantly down-regulated from -2,3 to -1,1 compared to Water + *B. cinerea*. Other notable genes with strong priming include those with proteolysis activity, with a range between -3 to -1,7 fold repressed. Other genes with repressed expression belong to auxin hormones and one to the ABA receptor (ABAPYL4). Furthermore, two genes of the little-known LATERAL ORGAN BOUNDARIES (LOB) were identified as repressed. Additional transcripts were functionally unassigned within the list.

Among the 57 differentially up-regulated genes (Table S2), there was one transcript encoding peroxidase activity with 2 fold increase compared to Water + *B. cinerea*, nine transcripts with protein kinase activity with between +1,1 to +2,1 fold, five transcripts with transcription regulatory activity, including SlMYB20, SLWRKY51 and SlWRKY72. Additional transcripts were functionally unassigned within the list.

Importantly, uncharacterised genes also show primed expression patterns. Of these, Avr9/Cf-9 rapidly elicited protein 75 (ACRE75; Solyc11g010250.1) was up-regulated 1,6-fold in Chitosan + *B. cinerea* in comparison to water + *B. cinerea* at 6 hpi (Table S2). ACRE genes have been previously studied and characterised as important genes involved in R gene-mediated and ROS gene-independent early plant defence responses (Durrant et al., 2000) and in response to methyl-jasmonate (MeJA) treatment (van den Burg et al., 2008). ACRE75 molecular functions are still to be deciphered and therefore research into its role and other members of the ACRE gene family in priming of chitosan was pursued.

### Role of ACRE genes in induced resistance against *Botrytis cinerea*

In order to investigate whether other members of the ACRE gene family display a similar priming profile to *ACRE75*, correlation analysis was performed on the subset of genes differentially expressed at 6 hpi. Genes with statistically significant similar profiles were identified (Table S3), which included *ACRE180* at a confidence value of 0.956. In addition, analysis of the samples later in the experiment, confirmed that both *ACRE75* and *ACRE180* are primed also at later time points (Fig S4).

In order to investigate whether primed expression of *ACRE75* and *ACRE180* genes may be involved in enhanced disease resistance, genes from *S. lycopersium* and ortholog genes in *N. benthamiana* were overexpressed using both transient and stable systems. For *SlACRE75*, best match against *N. benthamiana* genome was Niben101Scf03108g12002.1 (termed *NbACRE75*), sharing a 77.5% protein identity; (ii) For *SlACRE180*, the best match against the *N. benthamiana* genome was Niben101Scf12017g01005.1 (termed NbACRE180), with 49.5% protein identity. Arabidopsis ortholog analysis failed to identify hits for *ACRE75* and *ACRE1280* candidate genes. Constructs were produced with a fused GFP protein in the N-terminus and protein integrity was confirmed via Western blot. Proteins extracted from *N. benthamiana* leaves 48h after agro-infiltration and Western blot analysis confirmed that they were the expected sizes (Fig S5). Subcellular location of proteins was analysed via confocal microscopy of GFP fluorescence (Fig S6). Overexpression constructs were co-infiltrated with RFP-marker pFlub vector (McLellan et al., 2013) (Fig S6a) into *N. benthamiana* reporter lines CB157 (nucleus mRFP marker - Fig S6b) and CB172 (ER mRFP marker - Fig S6c). Free GFP accumulated in both cytoplasm and nucleus (Fig S6d), whereas GFP-SlACRE75 and GFP-NbACRE75 fusions accumulated exclusively in the nucleus and nucleolus of *N. benthamiana* cells (Fig S6e,f). Furthermore, GFP-SlACRE180 fusion accumulated exclusively in ER (Fig S6g), whereas GFP-NbACRE180 fusion accumulation was exclusively in peroxisomes (Fig S6h).

To further investigate the impact of overexpression of *ACRE* genes in disease resistance, the 4 constructs containing GFP-SlACRE75, GFP-SlACRE180, GFP-NbACRE75 and GFP-NbACRE180, and GFP-empty vector (EV), were agro-infiltrated into leaves of *N. benthamiana* plants, which were subsequently challenged with *B. cinerea*. Chitosan-induced resistance against *B. cinerea* was proven effective in *N. bethamiana* (Fig 4a). All GFP-SlACRE75, GFP-SlACRE180, GFP-NbACRE75 and GFP-NbACRE180-infiltrated *N. benthamiana* leaves showed a significant decreased in *B. cinerea* necrotic lesion size compared with the EV control (Fig 4b). To further analyse *ACRE75* and *ACRE180* biological functions and to confirm their role in plant resistance against *B. cinerea*, Arabidopsis plants were transformed to constitutively overexpress GFP-SlACRE75, GFP-SlACRE180, GFP-NbACRE75 and GFP-NbACRE180 proteins. Homozygous lines were identified and growth phenotype of transgenic plants was analysed by measuring rosette perimeter. No statistically significant differences were identified (Fig S7). Five-week-old plants were infected with *B. cinerea* and disease was scored at 6 dpi. Transgenic GFP-SlACRE75, GFP-SlACRE180, GFP-NbACRE180 and GFP-NbACRE75 overexpression plants all showed an enhanced resistance phenotype and significantly decreased *B. cinerea* lesion sizes in comparison to Col-0 and GFP-EV controls (Fig 4c). Furthermore, GFP-SlACRE75 and its homolog GFP-NbACRE75-overexpression plants showed a stronger resistance to *B. cinerea* than GFP-SlACRE180 and GFP-NbACRE180 overexpression lines at 6 dpi (Fig 4c).

**Figure 4.**
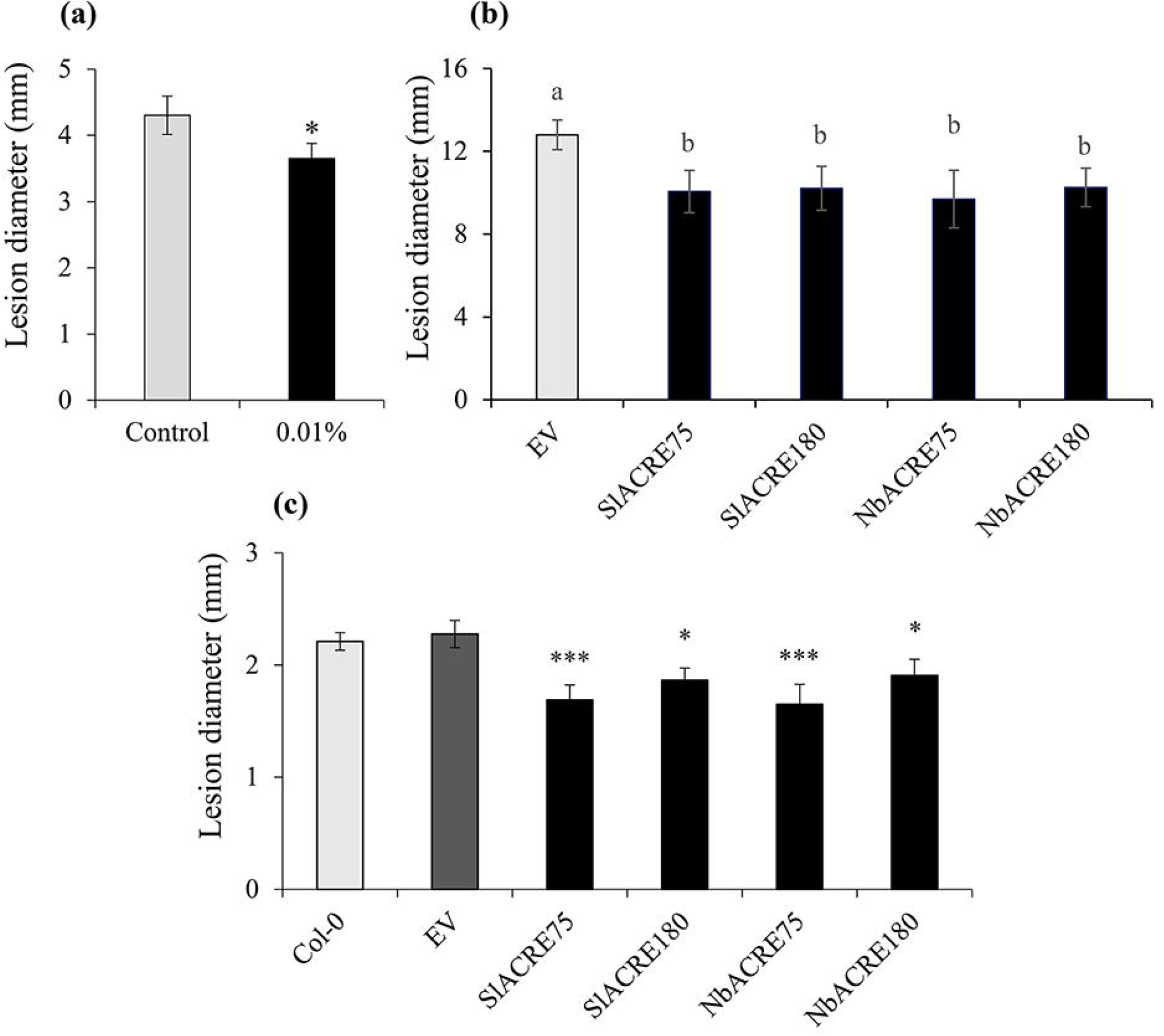
Functional characterisation of ACRE genes. **(a)** Chitosan-induced resistance in *Nicotiana benthamiana*. Disease lesions at 2 days post inoculation (dpi). Values represent means ± SEM (n=18). Asterisk indicates statistically significant differences between treatments (Student’s T. test, α=0.05). **(b)** Transient expression of constitutively active SlACRE75, SlACRE180, NbACRE75 and NbACRE180 in *N. benthamiana* against *B. cinerea*. Lesion size measurements were performed at 4 days post-infection (dpi). Values presented are means ± SEM (n=6). Different letters indicate statistically significant differences (ANOVA p < 0.05 followed by Tukey’s Post-hoc at 4 dpi). **(c)** *A. thaliana* transformed overexpression stable SlACRE75, SlACRE180, NbACRE75 and NbACRE180 infected with *B. cinerea*. Lesion sizes were measured at 6 days after inoculation (dpi). Values presented are means ± SEM (n=8-16). Asterisks indicate statistically significant differences (* p < 0.05, ** p < 0.01; *** p < 0.001).

## Discussion

We have assessed the capacity of chitosan to induce resistance against *B. cinerea* in different plant species and have linked its effect with priming of defence mechanisms. We have identified a formulation of chitosan that unlike some other formulations, can be easily dissolved in water and does not require infiltration. This opens possibilities to identify early-acting priming mechanisms in elicitor-induced resistance. Moreover, it enables opportunities for upscaling the use of chitosan as an elicitor of resistance in large-scale experiments due to the high-throughput nature of spraying the elicitor onto plants.

Treatments with chitosan resulted in induced resistance in *S. lycopersicum* (Fig 1a), *S. melongena* (Fig S1), Arabidopsis (Fig 1b) and *N. benthamiana* (Fig 4a) at a range of concentrations, which indicates that there are similar defence mechanisms acting in the response to fungal PAMPs. Moreover, treatments with chitosan resulted in the activation of basal resistance processes such as the deposition of callose at the cell wall (Fig 1c and d), which is considered an important factor for penetration resistance against invading pathogens (Oide et al., 2013). Expression of resistance was dependent on the concentration of chitosan used in Arabidopsis. In contrast, in tomato and aubergine the levels of resistance did not depend on the chitosan concentration. Moreover, chitosan-induced callose deposition in tomato and Arabidopsis did not follow a classical dose-response curve and the most effective treatments that activated callose were the lower concentrations of the elicitor (Fig 1c and d). This is likely to be dependent on the antimicrobial effects of chitosan (Fig S2) at higher concentrations. Other elicitors have been shown to trigger induced resistance phenomena at lower concentrations. For example, meJA treatment results in more effective protection against the pathogen *Fusarium oxysporum* f.sp. *lycopersici* when applied at lower concentrations (Król et al., 2015). In contrast, high doses of MeJA had detrimental effects on physiological processes and overall decreased protection efficiency. This, together with the observation that low concentrations of chitosan do not directly impact pathogen growth (Fig S2) suggests that there is a concentration threshold in the effect of chitosan-induced resistance.

Foliar applications of chitosan have been widely used to control disease development caused by numerous pests and pathogens (El Hadrami et al. 2010). However, few studies have investigated the role of chitosan as a priming agent and most have focused on its use as a seed priming elicitor mainly to improve germination and yield (Guan et al., 2009, Hameed et al., 2013). Here, we show that chitosan-induced resistance is based on priming of defence mechanisms. Our experiments confirmed that chitosan-induced resistance is not associated with growth reduction (Fig S3a), was durable and maintained for at least two weeks after treatment (Fig 2a), and that is based on a stronger accumulation of callose at the site of attackand accumulation of JA (Fig 2d) and JA-ile (Fig S3b). These results demonstrate that fungal growth arrest after chitosan treatment is not directly mediated by the toxicity effect of the chemical, as the infected leaves were formed after treatment and therefore were not sprayed with the elicitor. Moreover, these results demonstrate similar priming mechanisms after chitosan treatment to other elicitors, including Hx, which has been linked with priming of callose and JA against *B. cinerea* (Fernández-Crespo et al., 2017, Wang et al., 2014). Interestingly, however, despite many reported antagonistic and other crosstalk interactions between plant hormones (Robert-Seilaniantz et al., 2011), the concentrations of other plant hormones, SA and ABA, were not affected. This suggests that priming by chitosan does not result in the downregulation of other hormone-dependent signalling pathways, thereby maintaining an effective resistance status against other stresses.

In order to further explore priming of defence and to unravel the transcriptional mechanisms behind chitosan-induced resistance, we performed transcriptome analysis. In our experiment, using a concentration of chitosan that is associated with priming but with no direct antimicrobial effect, we identified early-acting differential transcriptomic changes. Results demonstrate that chitosan treatments do not result in major transcriptional changes (Fig 3a). In contrast, comparison of treatment against Water + Mock revealed and Chitosan + *B. cinerea* shows a higher number of DEGs (Fig 3b and c), thus responding to the priming nature of the elicitor in the first instance.

Panther enrichment analysis showed that at 6hpi, the number of down-regulated DEGS was more than three times up-regulated ones for Chitosan + *B. cinerea* (203 down-regulated and 57 DEGs up-regulated). This suggests that tomato plants might repress susceptible factors in order to reduce *B. cinerea* manipulation of host defences (El Oirdi et al., 2011, Temme and Tudzynski, 2009). Interestingly, some of the down-regulated transcripts have cysteine-type peptidase activity (Table S1). These proteins have been reported to have a role in immunity against pathogens including *B*.*cinerea* (Pogány et al., 2015). Other down-regulated genes are related to plant hormone activity; including ethylene AP2/ERF transcription factors and ABA PYL receptors (SlABAPYL4), reported to be involved in defence responses, which act as positive or negative regulators of JA/ET-dependent defences against *B. cinerea* (Cantu et al., 2009, Moffat et al., 2012). Up-regulated genes included transcripts with peroxidase and transcription regulatory activity, such as peroxidase 5, SlMYB20, SLWRKY51 and SlWRKY72, CONSTANS-like protein with zinc finger binding domain and NAC domain protein and a RING-type E3 ubiquitin transferase involved in protein degradation. These genes have been linked with defence responses (Serrano et al., 2018), which could be result in priming of the tomato immune system against *B. cinerea* infection.

Transcriptomic (Table S2, Fig S4) and qRT-PCR (Fig S4) analyses showed that chitosan can prime *ACRE75* for a faster and stronger expression after infection with *B. cinerea*. ACRE genes have been linked to plant defence responses. Similar genes were previously identified in tobacco cells to exhibit rapid Cf-9–dependent change in expression through gene-for-gene interaction between the biotroph pathogen *Cladosporium fulvum* avirulence gene (Avr9) and tomato resistance Cf-9 gene (Durrant et al., 2000). To determine the role of ACRE genes in priming by chitosan, we searched for other ACRE genes showing similar expression profiles to *ACRE75* and this revealed that *ACRE180* displays a similar priming profile. This was more evident at 9 hpi (Fig S4) than at 6 hpi, suggesting that the role of ACRE180 is later time than ACRE75. Subcellular localisation may indicate why priming of these genes does not occur at the same time; whereas ACRE75 accumulates exclusively in the nucleus and nucleolus (Figure S6e and f), ACRE180 accrues in the ER and peroxisomes (Figure S6g and h). This suggests different molecular functions of these proteins as they tag different cell organelles. Moreover, it could be plausible that ACRE75 and ACRE180 are part of the same signalling pathway, one working upstream of the other, therefore justifying the delayed transcription and activity of ACRE180.

The roles of ACRE75 and ACRE180 in chitosan-induced priming were investigated by overexpressing these genes in transient and stable systems, in *N. benthamiana* and Arabidopsis, respectively. Moreover, we aimed to identify any *N. benthamiana* and Arabidopsis ACRE75 and ACRE180 analogues. BLAST analysis of tomato ACRE75 identified a very low amino acid identity sequence (39%) and ACRE180 failed to identify any Arabidopsis homologue. In contrast, *N. benthamiana* ACRE75 and ACRE180 homologues were putatively identified. ACRE75 and ACRE180 lack signal peptides, which suggests they might encode small proteins involved in signalling or antimicrobial activity within the infected cell. Similar to the exclusive production of glucosinolates compounds in Brassica plants (Matthaus & luftmann 2000) it is likely that ACRE75 and ACRE180 are involved in the production of unique compounds to Solanaceae plants. Overexpression of *SlACRE75* and *SlACRE180*, and their *N. benthamiana* orthologues results in induced resistance against *B. cinerea* (Fig 4b and c). Therefore, our results confirm involvement of ACRE genes in plant immunity and suggest an involvement in chitosan-induced priming due to their expression profiles. Interestingly, the induced resistance effect was greater in Arabidopsis plants overexpressing ACRE75 in comparison to ACRE180 (Fig 4c), which could corroborate our evidence of earlier activity of ACRE75, therefore being more effective during early resistance response. More work in needed to unravel the molecular function of ACRE75 and ACRE180 in the expression of priming mechanisms. Nevertheless, fine-tuning of priming-based mechanisms under the control of *SlACRE75, SlACRE180, NbACRE75* and *NbACRE180* could facilitate its incorporation into other crop species for the enhancement of cross tolerance to old and emergent pest and pathogens, and other challenges. The results unveiled potential molecular pathways involved in chitosan-induced priming of resistance in tomato against *B. cinerea*, potentially applicable to other crops.

## Acknowledgement

The authors thank Prof Nicola Stanley-Wall for the academic supervision, Prof Murray Grant for his assistance during HPLC analysis of plant defence hormones, Prof Jurriaan Ton for initial advice on the project, laboratory access, resources, support and intellectual input, Dr Katherine Wright for her support during the confocal microscopy analysis, Dr Colin Alexander for his help with the statistics described in the paper and Dr Mike Roberts for providing spores of *B. cinerea* and extremely valuable input with the microarray analysis. We thank ChiPro GmbH for providing the commercial formulation of chitosan. We are grateful for financial support from the Rural and Environmental Science and Analytical Services (RESAS) Division of the Scottish Government (2011-16) under its Environmental Change and Food, Land and People Research Programmes. This work is funded by AHDB (HDC) Studentship CP105 was secured by A.N and N.H to support D.DV doctoral programme, and the BBSRC Future Leader Fellowship BB/P00556X/1 and BB/P00556X/2 to E.L.

## Author contribution

All bioassays were performed by D.DV and E.L. Transcriptome analysis was done by J.M and P.E.H Data analysis was performed by D.DV, N.H, P.E.H and E.L. Intellectual input was provided by D.DV, N.H, P.E.H, E.L and A.N. Project was conceived and supervised by N.H and A.N. The manuscript was written by D.DV and E.L with input from all authors.

## Figure Legends

**Figure S1. Chitosan-induced resistance in *Solanaceae melongena* (aubergine)**. Disease lesions at 3 dpi. Values represent means ± SEM (n=10). Different letters indicate statistically significant differences among treatments (Least Significant Differences, α=0.05).

**Figure S2. Chitosan and Switch fungicide antifungal activity against *Botrytis cinerea***. Bars represent means of fungal growth diameter (± SEM, n=5) at 4 days after inoculating PDA-containing Petri dishes with 5 mm agar plugs of actively growing *B. cinerea* mycelia. Different letters indicate statistically significant differences among treatments (Least Significant Differences, α=0.05)

**Figure S3. Chitosan-induced resistance is based on priming. (a)** Relative Growth Rate (RGR) per week of tomato plants 1 and 2 weeks after treatment with 0.01% chitosan. Values represent means ± SEM (n=10). Asterisk indicates statistically significant differences among treatments (Student’s T. test, α=0.05) **(b)** Mass-spectrometry quantification (% of peak area) of Jasmonic acid-isoleucine (JA-ile) at 24h post inoculation. Values represent means ± SEM (n=4). Different letters indicate statistically significant differences among treatments (Least Significant Differences, α=0.05)

**Figure S4. Validation of microarray expression results**. Expression profile obtained in the microarray (a) and in the analysis by RT-q-PCR (b) of a subset of 9 genes at 9 hpi with *Botrytis cinerea*

**Figure S5. Western Blot analysis**. Expression of proteins by immunoblot analysis of GFP-SlACRE75, GFP-SlACRE180, GFP-NbACRE75 and GFP-NbACRE180 fusion proteins in *N. benthamiana* leaves at 48 h after agroinfiltration. Expected protein sizes were (i) SlACRE75= 14.79 + 26 KDa GFP= 40.8 KDa; (ii) SlACRE180= 10.86 +26= 36.8 KDa; (iii) NbACRE180= 11.74+26= 37.7 KDa; and (iv) NbACRE75= 14.6+26= 40.7 Kda. Proteins were separated by SDS–PAGE and analysed by immunoblotting. A GFP-specific antibody was used for detection of GFP-fusion protein. Equal loading of total proteins was examined by Ponceau staining (PS). Three lanes represent 3 replicates per construct GFP-SlACRE75, GFP-SlACRE180, GFP-NbACRE75, GFP-NbACRE180, and a GFP-non-protein/ empty vector (control).

**Figure S6. Subcellular location of ACRE proteins**. Confocal microscopy observation of **(a)** pFlub vector as a RFP-peroxisome tagged marker, **(b)** nucleus mRFP marker, **(c)** ER mRFP marker, **(d)** free GFP in cytoplasm and the nucleus, **(e)** GFP-SlACRE75 and **(f)** GFP-NbACRE75 fusions in the nucleus and nucleolus, **(g)** GFP-SlACRE180 fusion in the ER and **(h)** GFP-NbACRE180 fusion in the peroxisomes.

**Figure S7. Growth analysis**. Perimeter in cm of rosettes from Arabidopsis lines overexpressing GFP-Empty vector (EV), GFP-SlACRE75, GFP-SlACRE180, GFP-NbACRE75 and GFP-NbACRE180 constructs. Values represent means ± SEM (n=8-16). n.s = not significant differences between treatments (One-way ANOVA, α=0.05)

## Notes

https://earray.chem.agilent.com/earray

