## Supplementary figures and images for "Chitosan primes plant defence mechanisms against *Botrytis cinerea*, including expression of Avr9/Cf-9 rapidly-elicited genes"

### Figure S1

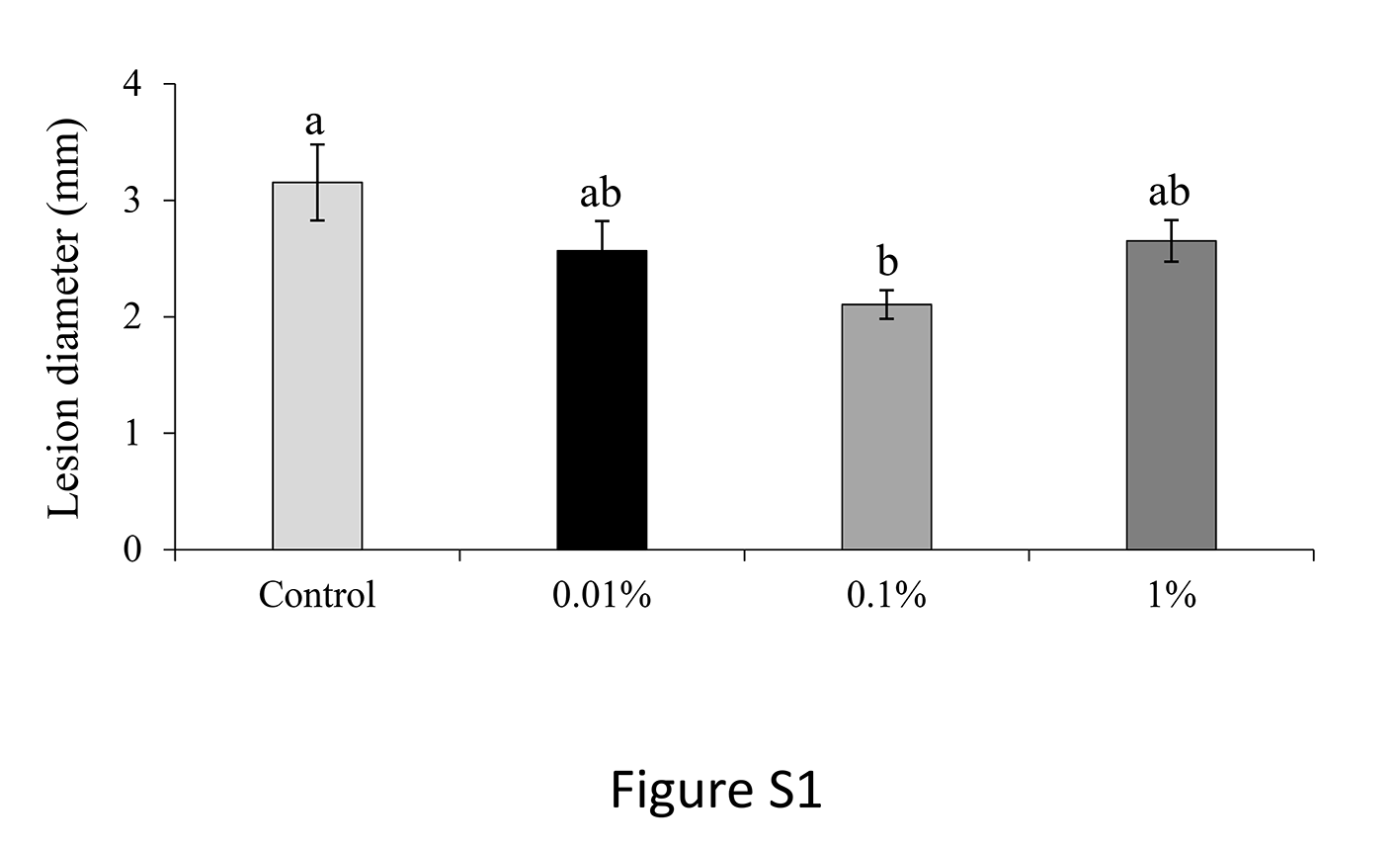

### Figure S2

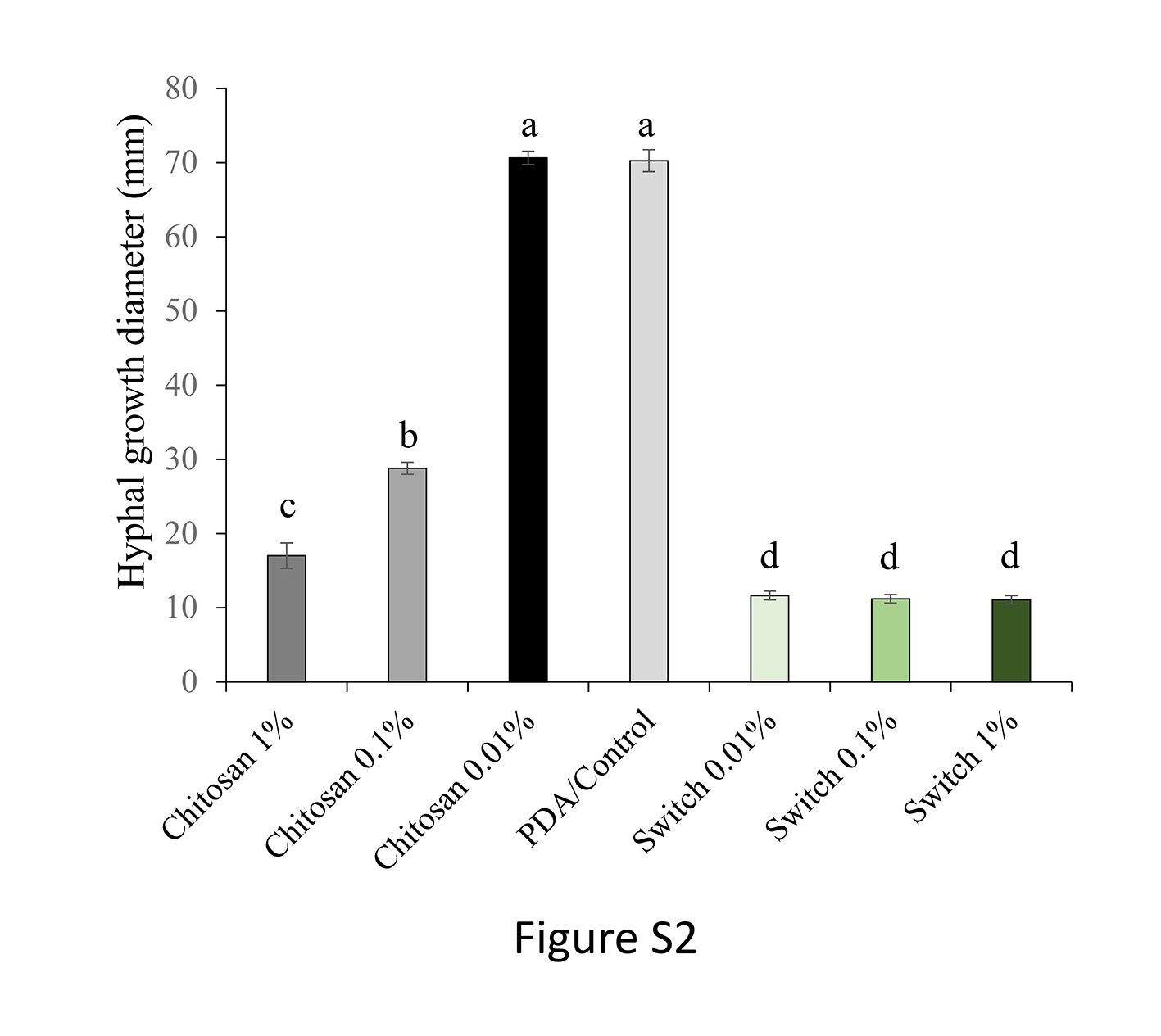

### Figure S3

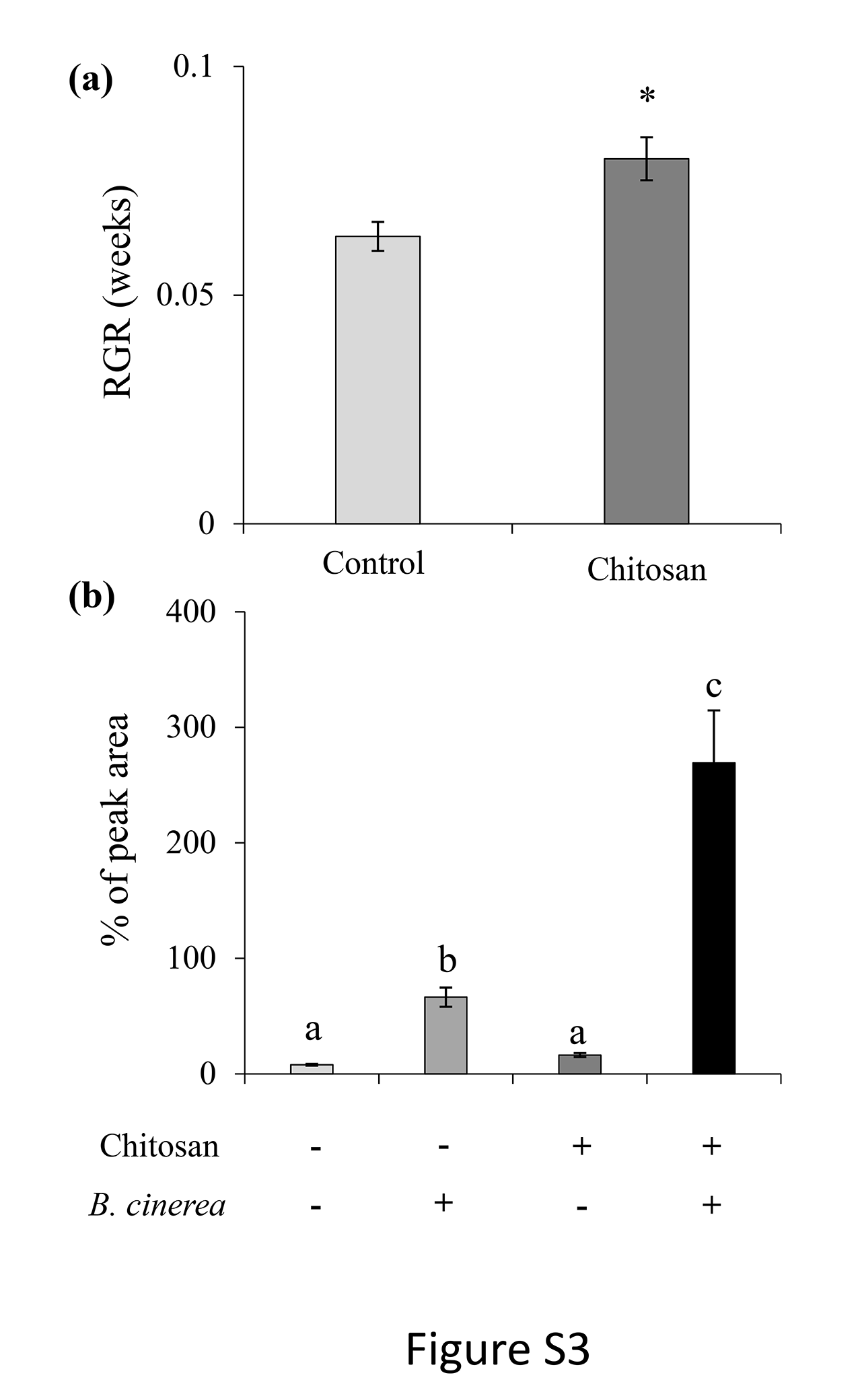

### Figure S4

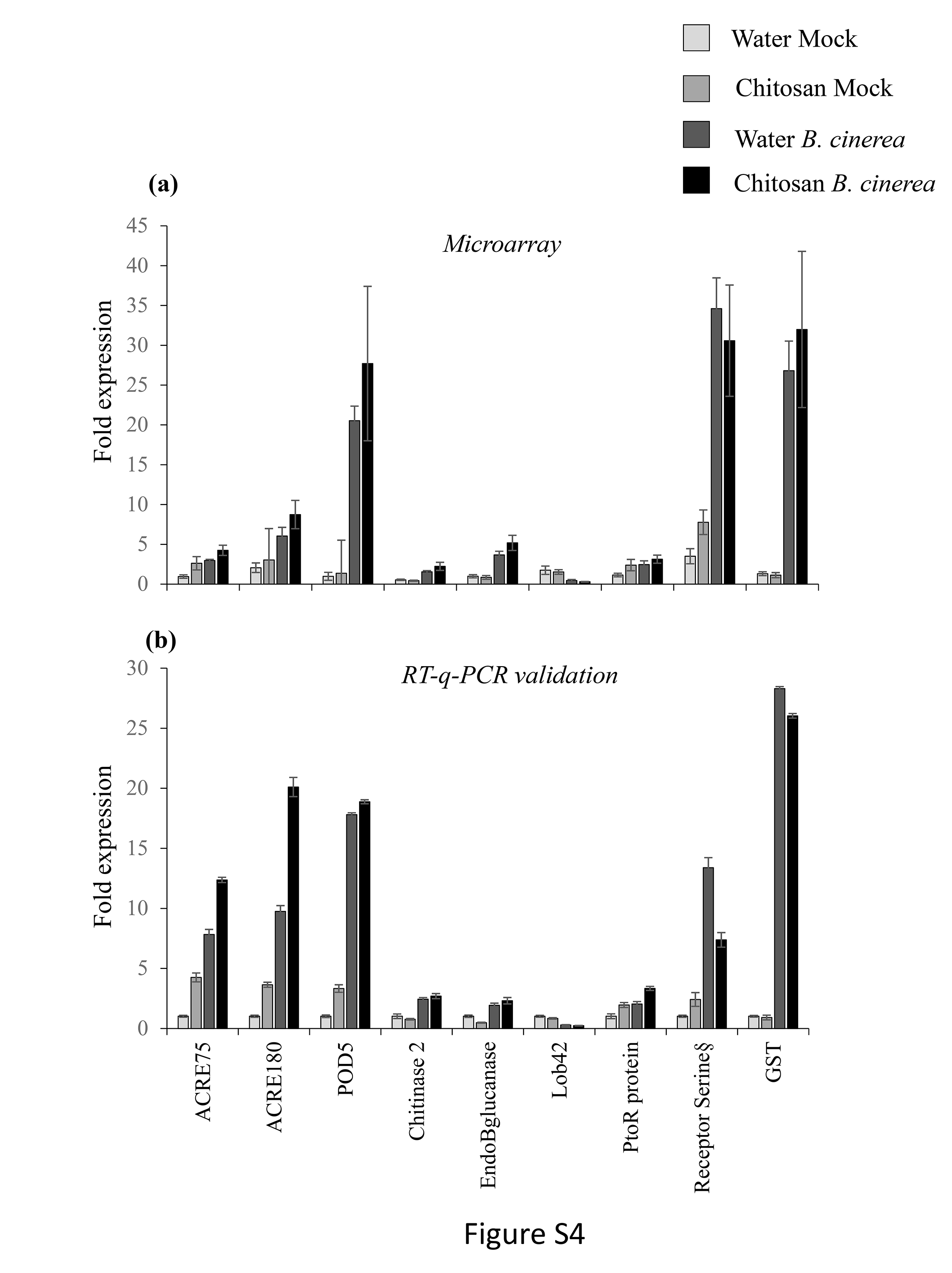

### Figure S5

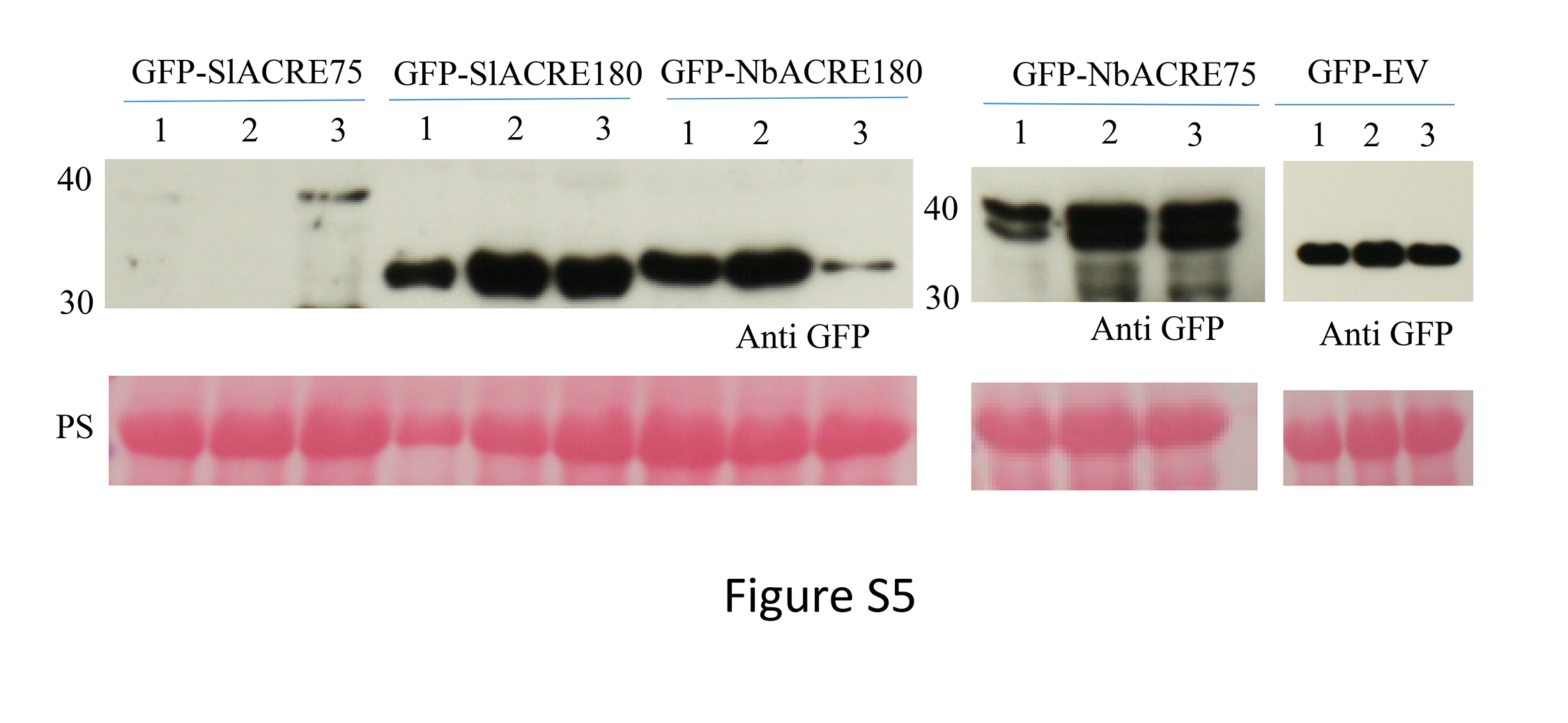

### Figure S6

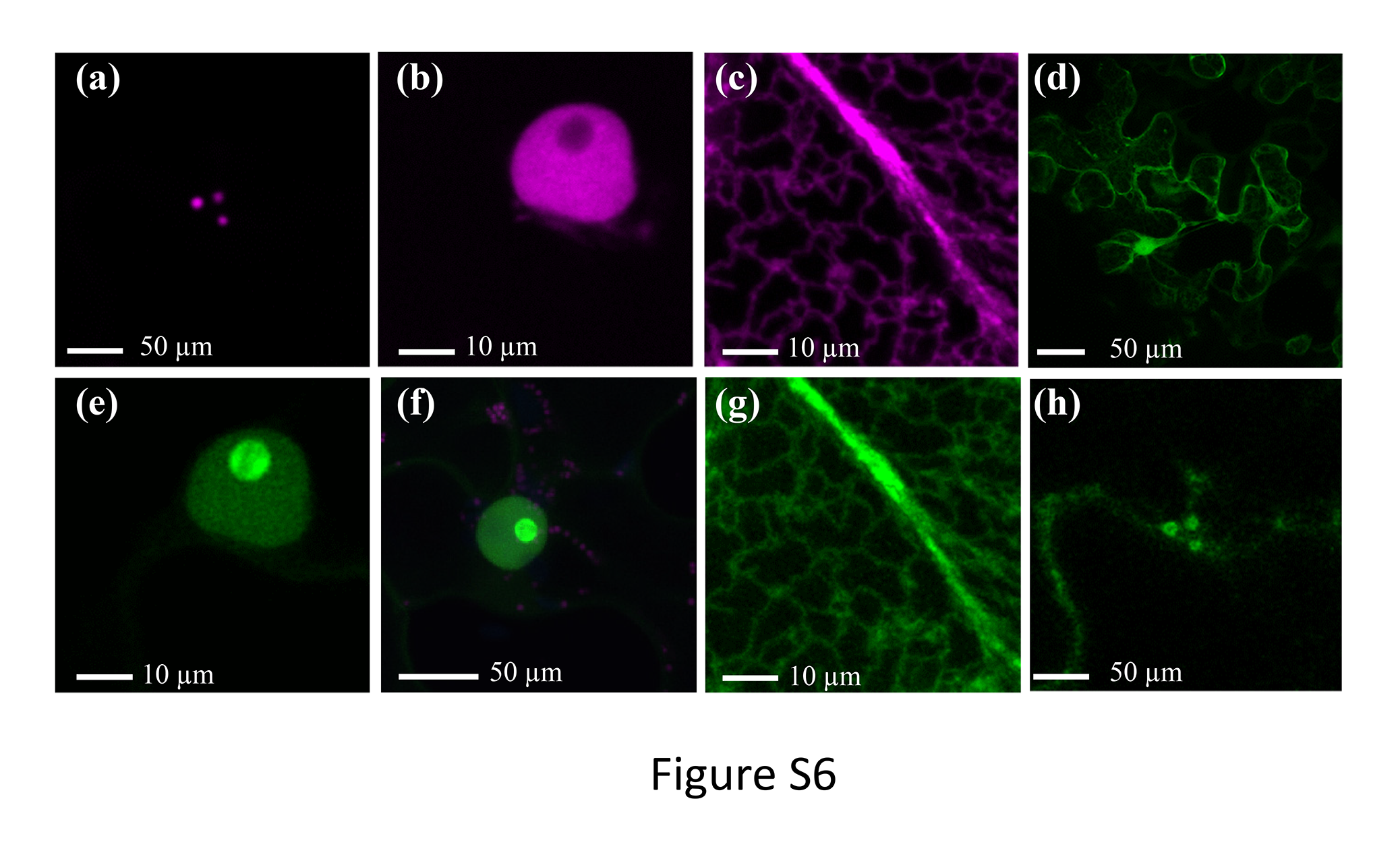

### Figure S7

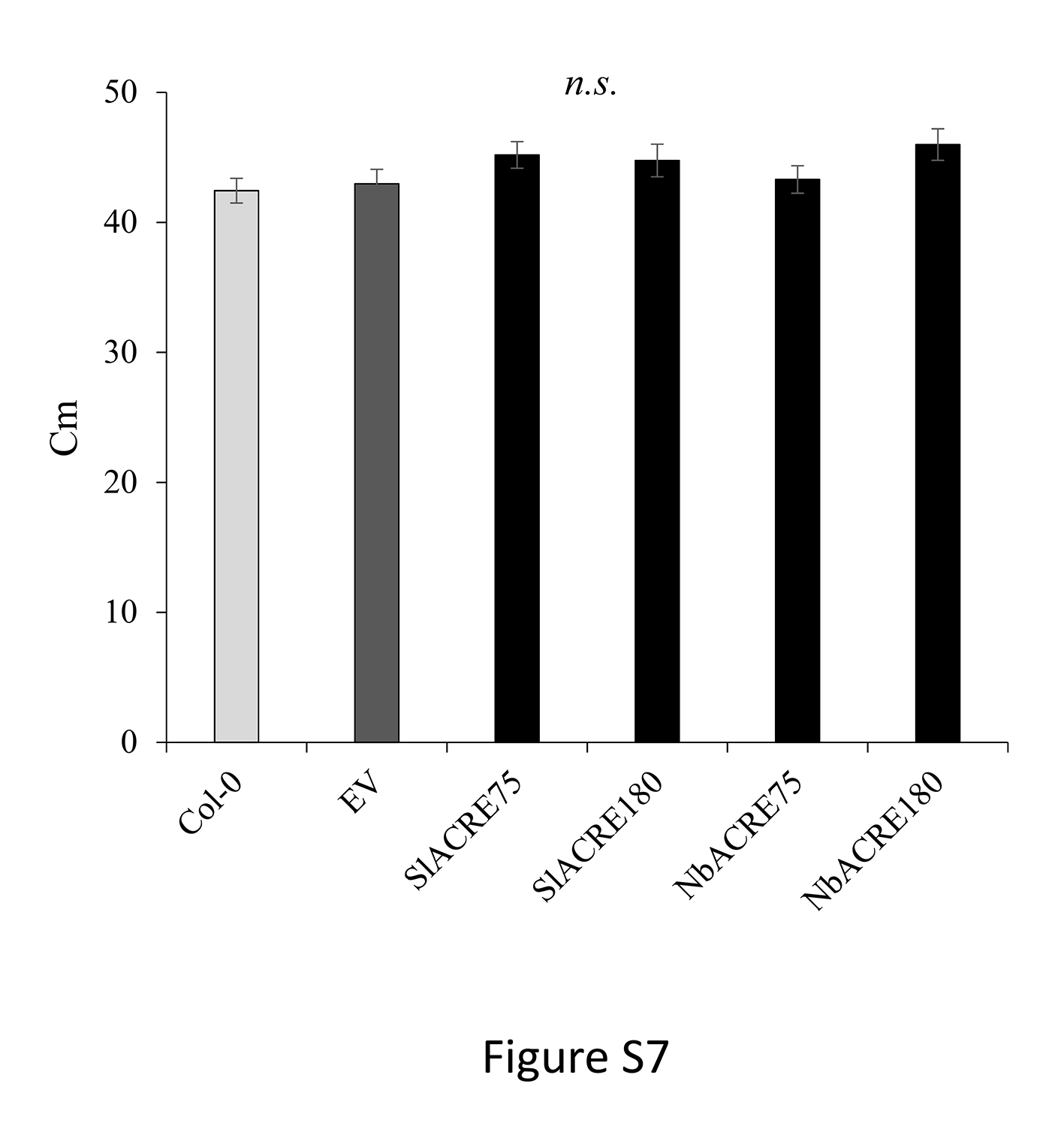
